# *FLOWERING LOCUS C* drives delayed flowering in *Arabidopsis* grown and selected at elevated CO_2_

**DOI:** 10.1101/2023.06.15.545149

**Authors:** Aleah Henderson-Carter, Hannah Kinmonth-Schultz, Lena Hileman, Joy K. Ward

**Affiliations:** Department of Integrative Biology, University of Texas at Austin, Austin, TX 78712, USA; Department of Biology, Tennessee Technological University, Cookville, TN 38505, USA; Department of Ecology and Evolutionary Biology, University of Kansas, Lawrence, KS 66045, USA; College of Arts and Sciences, Department of Biology, Case Western Reserve University, Cleveland, OH 44106, USA

**Keywords:** *Arabidopsis*, elevated [CO_2_], *FLOWERING LOCUS C* (*FLC*), *FLOWERING LOCUS T* (*FT*), flowering time, global change, *LEAFY (LFY)*, *SUPPRESSOR OF CONSTANS 1* (*SOC1*), selected genotype (SG), vernalization

## Abstract

- Altered flowering time at elevated [CO_2_] is well documented, although mechanisms are not well understood. An *Arabidopsis* genotype previously selected for high fitness at elevated [CO_2_] (SG) showed delayed flowering and larger size at flowering when grown at elevated (700 ppm) versus current (380 ppm) [CO_2_]. This response was correlated with prolonged expression of *FLOWERING LOCUS C* (*FLC*), a vernalization-responsive floral repressor gene.
- To determine if *FLC* directly delays flowering at elevated [CO_2_] in SG, we used vernalization (extended cold) to downregulate *FLC* expression. We hypothesized that vernalization would eliminate delayed flowering at elevated [CO_2_] through the direct reduction of *FLC* expression, eliminating differences in flowering time between current and elevated [CO_2_].
- We found that with downregulation of *FLC* expression via vernalization, SG plants grown at elevated [CO_2_] no longer delayed flowering compared to current [CO_2_]. Thus, vernalization returned the earlier flowering phenotype, counteracting effects of elevated [CO_2_] on flowering.
- This study indicates that elevated [CO_2_] can delay flowering directly through *FLC*, and downregulation of *FLC* under elevated [CO_2_] reverses this effect. Moreover, this study demonstrates that increasing [CO_2_] may potentially drive major changes in development through *FLC*.

## Introduction

A critical aim in the ecology and evolutionary biology field is to better understand the mechanisms driving changes in plant phenology events (Gamage *et al*., 2018). Plant phenology is the study of life cycle changes, such as leaf-out and flowering time, and their fundamental role in terrestrial ecosystems. Flowering time is an important phenology event because changes in the timing of the transition from the vegetative to reproductive phase can impact development, plant-pollinator interactions, fruiting, and seed-set Fitter & Fitter, 2002; Cleland *et al*., 2007). Over the last several decades, plant biologists have been investigating the effects of anthropogenic global change on flowering time. Classic hallmarks of anthropogenic global change have included rising global temperatures, loss in biodiversity, land-use change, and rising atmospheric CO_2_ concentrations (Gamage *et al*., 2018). The immediate effects of rising [CO_2_] on physiological and ecosystem functioning have been explored; however, the molecular mechanisms driving flowering time shifts in response to rising [CO_2_] remains unclear.

Flowering time affects productivity, plant-pollinator interactions, and fitness (Fitter & Fitter, 2002; Cleland *et al*., 2007). Global change factors have major impacts on flowering time, with the effects of changing temperature receiving the most attention (Becklin *et al*., 2016; Tun *et al*., 2021). An examination of the warming effects on flowering time concluded that warming alone under-predicts flowering time change (Wolkovich *et al*., 2012). Therefore, although temperature is a key driver of flowering time, additional global change determinants of inter- and intraspecific flowering time variation remain unexplained (Jentsch *et al*., 2009; Tun *et al*., 2021). Specifically, the direct effects of rising atmospheric CO_2_ concentration ([CO_2_]), which affect plant physiology and development (Gamage *et al*., 2018), must be considered as a key driver of altered flowering time responses under global change scenarios.

Atmospheric [CO_2_] (>400 ppm) is predicted to increase to *c*. 800 ppm by 2100 (Allen *et al*., 2018). Elevated [CO_2_] increases plant growth rates and alters development time in well over half of wild and crop species (Springer & Ward, 2007). If these projections hold, under the assumption that international course-correction efforts fail, plant life, especially for C3 photosynthetic pathway species, may not maintain physiological and developmental processes in this high [CO2] world. Studies suggest that elevated [CO_2_] influences flowering through increased photosynthesis, leading to more carbon resources for developmental processes (Gamage *et al*., 2018). However, the direction and magnitude of flowering time shifts under altered [CO_2_] are highly variable, including both accelerations and delays, which are not always predictable by accelerated growth rates (Morison & Lawlor, 1999; Ward & Kelly, 2004; Springer & Ward, 2007). To predict how plants will respond to global change, we need to better understand [CO_2_] effects on genetic flowering pathways and subsequent effects on downstream impacts on visible flowering time.

Flowering time pathways integrate environmental and endogenous cues that converge to regulate the expression of key floral genes via multiple signaling pathways (Conti, 2017). The MADS-box transcription factor *FLOWERING LOCUS C* (*FLC*), a major floral repressor of flowering in *Arabidopsis*, may be especially important for understanding [CO_2_]-driven flowering time changes (Springer *et al*., 2008). *FLC* represses flowering in winter-annual and perennial Brassicaceae species until it is repressed by prolonged cold (vernalization) to facilitate spring flowering (Bratzel & Turck, 2015).

An *Arabidopsis* genotype selected over multiple generations at elevated [CO_2_] for increased fitness (Selected Genotype; SG) exhibited delayed flowering by 7 to 10 d, along with higher seed output when grown at 700 ppm relative to 380 ppm CO_2_ (Ward & Strain, 1997; Ward & Strain, 1999; Ward *et al*., 2000). This is in contrast to a closely related Control Genotype (CG) which exhibited similar flowering at both 380 ppm and 700 ppm CO_2_, providing a powerful comparative model (Ward *et al*., 2000; Springer *et al*., 2008). In SG, expression of *FLC* was elevated and extended across a greater period of plant development at 700 ppm [CO_2_], correlating with the delayed flowering phenotype. Altered expression of downstream floral initiator genes, *SUPPRESSOR OF OVEREXPRESSION OF CONSTANS 1* (*SOC1*) and *LEAFY* (*LFY*), in SG grown at 700 ppm were also correlated with delayed flowering. These alterations in gene expression were not observed in CG grown at 700 ppm CO_2_ (Springer *et al*., 2008), which was correlated with no effects of elevated [CO_2_] on flowering time. This highlights a candidate mechanism involving *FLC* that may be driving this delayed flowering at elevated [CO_2_].

However, it remains unknown whether elevated *FLC* expression in a high [CO_2_] environment is directly linked to the delayed flowering phenotype in *Arabidopsis*.

Many species do not initiate flowering until they have experienced vernalization which promotes flowering during the spring (Reeves *et al*., 2007). The *Arabidopsis* vernalization responsive gene, *FLC*, inhibits flowering by repressing key floral initiator genes, *FLOWERING LOCUS T* (*FT*) and *SOC1* (Searle *et al*., 2006; Conti, 2017). Together *FT* and *SOC1* positively regulate an additional key floral inducer, *LFY* (Lee & Lee, 2010). Vernalization leads to a series of epigenetic modifications at *FLC* that result in mitotically stable repression (Sung *et al*., 2006).

Sequence variation at the *FLC* locus modulates the requirement for vernalization-induced *FLC* repression (Shindo *et al*., 2006; Madrid *et al*., 2021). This variation is frequently responsible for natural flowering time variation in *Arabidopsis* and other Brassicaceae (Akter *et al*., 2018).

Vernalization was not necessary for SG or CG to flower (Springer *et al*., 2008). However, given the prolonged high expression of *FLC* under 700 ppm [CO_2_] in SG, this elevated [CO_2_]-adapted genotype may be vernalization-responsive at elevated but not current [CO_2_]. Thus, vernalization can be used as a tool to assess the role of *FLC* in CO_2_-induced delayed flowering.

Here, we determined if *FLC* expression drives shifts in flowering time under elevated [CO_2_]. To achieve this, we manipulated *FLC* expression through vernalization treatments in SG and CG under current and elevated [CO_2_]. We hypothesized that vernalized SG plants, with reduced *FLC* expression, would be rescued from elevated [CO_2_]-induced delays in flowering. Because *FLC cis* variation is responsible for natural variation in *Arabidopsis* flowering time, we also assessed *FLC* sequence variation between SG and CG. This study provides further insights on the causal mechanism driving flowering shifts under elevated [CO_2_].

## Materials and Methods

Soil-grown *Arabidopsis thaliana* (L.) Heynh SG and CG genotype (Ward *et al*., 2000) seeds were maintained at 400 and 800 ppm [CO_2_], 22:18 °C (day:night), and 14:10 h photoperiod at *c*. 650 μmol m^−2^ s^−1^ light intensity. At the development stage of 4-5 true leaves, a subset of randomly selected SG and CG were moved into vernalization (5 °C; 24-hour cycle) at 400 ppm or 800 ppm [CO_2_] for 30 d and then were returned to their prior growing conditions at the completion of this cold treatment. Rosettes were harvested for dry shoot weight and leaf number at flowering when inflorescences were 1 cm in height, which was used as a measurement to standardize time since growth conditions varied across treatments. In this way, relatively higher leaf number at flowering would represent a delay, whereas lower leaf number would represent an acceleration.

Whole rosettes were also harvested between dawn and noon at the start of vernalization (time 0), 2, 3, and 4 weeks after the start of vernalization, and 5 days post-vernalization. *FLC, FT, SOC1*, and *LEAFY* (*LFY)* transcript levels, normalized against *UBC9* (AT4G27960) and *YLS8* (AT5G08290), (Czechowski *et al*., 2005; Hong *et al*., 2010) (Primers in Table S1) were measured by RT-qPCR (Ito *et al*., 2012).

The *FLC* genomic sequence was pulled from SG and CG genomes assembled against the *Arabidopsis thaliana* reference Col-0 reference genome (TAIR version 10) (Rhee *et al*., 2003). Differences across genotypes, [CO_2_] and vernalization were assessed using ANOVA and Tukey post-hoc tests (R version 4.11) (see Supplemental Methods for additional details).

## Results

### Vernalization eliminated delayed flowering in SG grown at elevated [CO_2_]

To assess whether elevated [CO_2_] (800 ppm) acts through *FLC* to delay flowering in SG, we grew non [CO_2_]-responsive CG and [CO_2_]-responsive SG genotypes in 400 and 800 ppm [CO_2_] and treated them with vernalization to downregulate *FLC* (Sung & Amasino, 2004; Shindo *et al*., 2006; Sung *et al*., 2006). We hypothesized that SG grown at 800 ppm would flower earlier (as measured by leaf number at flowering) when vernalized compared to non-vernalized due to reduced *FLC* expression.

Flowering time significantly differed across genotypes, [CO_2_], and vernalization treatments (*p* < 0.0001). Interactive effects between [CO_2_] and vernalization on flowering time were observed in SG only (*p* < 0.0001). At 800 ppm [CO_2_], non-vernalized SG had significantly higher, and more variation in leaf number at flowering than vernalized SG (p < 0.0001; Fig. 1). Importantly, there was no significant difference in flowering time between non-vernalized and vernalized SG plants grown at 400 ppm [CO_2_] compared with vernalized SG grown at 800 ppm [CO_2_]. This indicated that vernalization eliminated SG’s delayed flowering at 800 ppm [CO_2_], leading to similar flowering time as SG grown at 400 ppm [CO_2_] (Fig. 1). In CG, we found similar flowering times across all [CO_2_] and vernalization treatments (Fig. 1).

**Figure 1.**
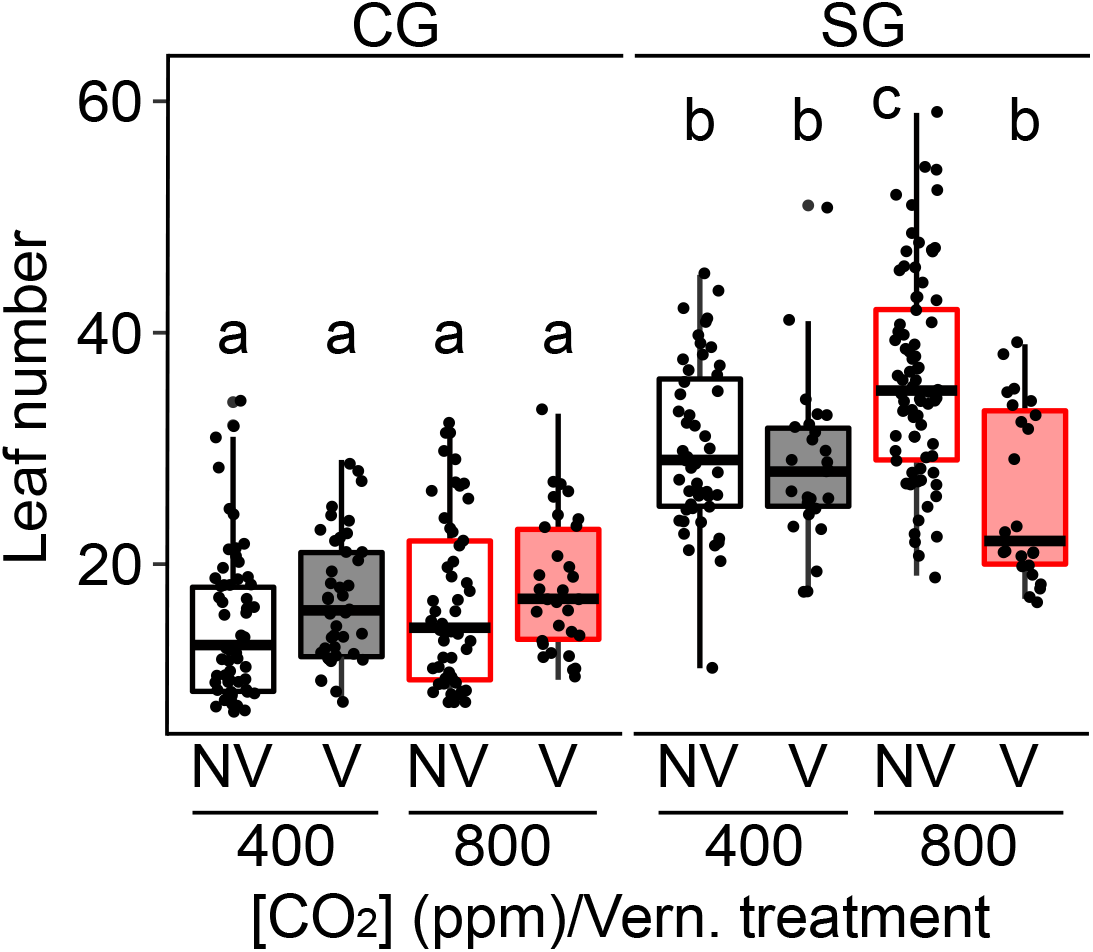
Flowering time measurements of *Arabidopsis thaliana* Control Genotype (CG) and Selected Genotype (SG) in non-vernalized (NV) and vernalized (V) conditions under elevated [CO_2_] (800ppm) and ‘current’ [CO_2_] (400ppm). Boxes indicated 25% and 75% quartiles and the heavy black line represents the median. Whiskers extend to highest value with 1.5 x the interquartile range. Lower-case letters indicate differences in significance. Each point represents a single individual.

Because plant size correlates with fitness and total seed number (Sletvold, 2002; Younginger *et al*., 2017), we assessed biomass at flowering. Consistent with leaf number at flowering, biomass varied significantly between genotypes, with higher biomass at flowering in SG relative to CG (p < 0.0001; Fig. S1). Generally, non-vernalized plants had higher biomass at bolt than vernalized plants (*p <* 0.0001; Fig. S1), although that was not the case for CG grown at 800 ppm [CO_2_] (Fig. S1). Notably, for SG plants grown at 800 ppm [CO_2_], vernalization resulted in *c*. 40% reduced biomass at flowering compared to non-vernalization (*p <* 0.0001; Fig. S1), and these plants also flowered at a lower leaf number as noted above. On the other hand, SG plants grown at 400 ppm CO_2_ flowered at the same leaf number whether exposed to vernalization or not (Fig. 1), even though non-vernalized plants had much higher biomass at flowering (Fig. S1). This indicates the independence of developmental timing as assessed by leaf number at flowering relative to biomass produced at flowering.

### Return to normal flowering time at 800 ppm CO_2_ is correlated with *FLC* downregulation in SG

To confirm *FLC* expression was repressed by vernalization at 800 ppm CO_2_ and to further assess downstream effects of [CO_2_] on flowering gene expression, we measured *FLC, FT, LFY*, and *SOC1* expression across all genotypes and treatments. We expected relatively high and more sustained expression of *FLC* in late-flowering, non-vernalized SG in 800 ppm [CO_2_] and relatively lower levels of *FLC* expression across all other early flowering treatments.

Additionally, we expected a negative correlation between *FLC* and downstream *FT, LFY*, and *SOC1* expression.

In CG, *FLC* declined in the early portions of the life cycle and was stably repressed across treatments regardless of vernalization (Fig. 2a), although expression at time zero was slightly elevated in 800 ppm [CO_2_] (*p* < 0.0001). Consistent with delayed flowering, *FLC* expression was significantly higher in non-vernalized compared to vernalized SG plants grown at 800 ppm [CO_2_] (*p* < 0.0001) across all developmental time points after time zero. There were no differences in *FLC* expression among vernalized SG in 800 ppm and vernalized/non-vernalized SG at 400 ppm [CO_2_] (Fig. 2b). Thus, vernalization of SG at 800 ppm [CO_2_] successfully downregulated *FLC*, supporting *FLC*’s role as a flowering repressor under elevated [CO_2_].

**Figure 2.**
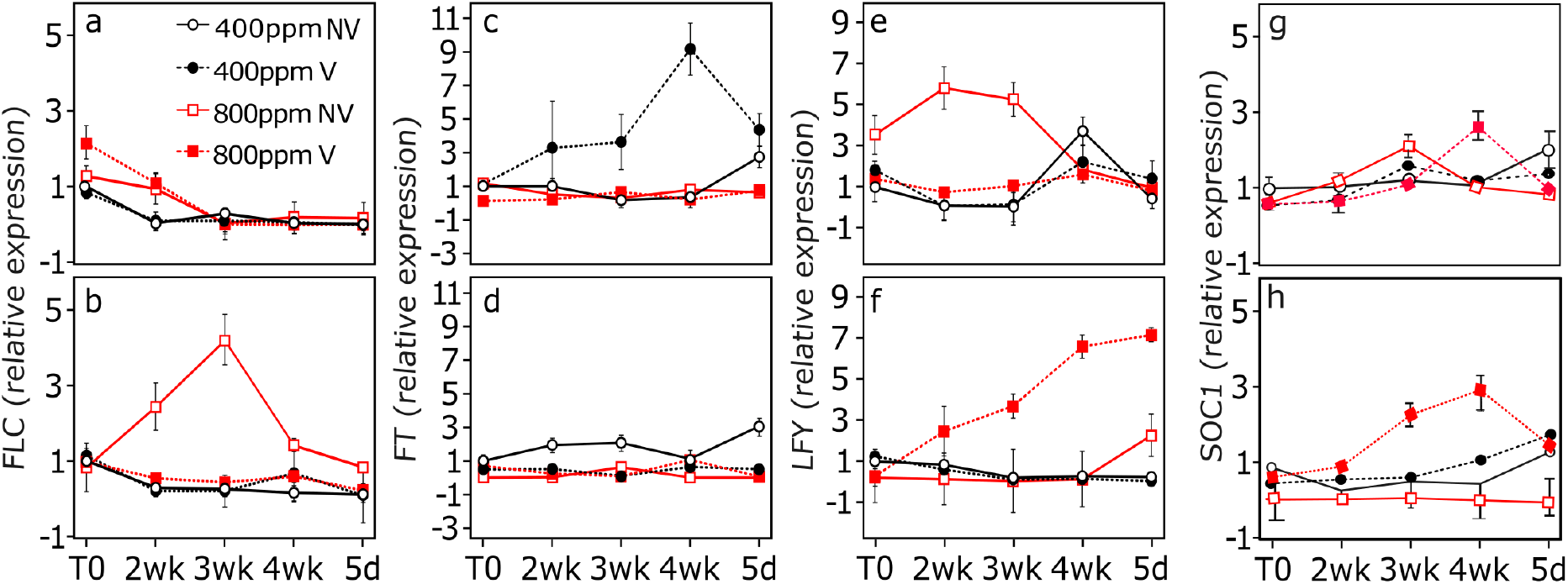
The expression patterns of *FLOWERING LOCUS C* (*FLC*), *FLOWERING LOCUS T* (*FT*), and *LEAFY* (*LFY*) genes vary across Selected Genotype (SG) and Control Genotype (CG). Expression of *FLC, FT, LFY*, and *SOC1* in CG (**a, c, e, g**) and SG (**b, d, f, h)** in plants grown at 400 and 800 ppm [CO_2_] and either not vernalized (NV) or vernalized for four weeks after plants reached four to five true leaves.

*FT, LFY*, and *SOC1* expression did not show a consistent pattern of negative correlation with *FLC* (Fig. 2c-g). *FT* expression was uniform across collection periods in CG 800 NV, 800V, and 400 NV. *FT* expression analyzed from CG 400 V deviated from all other CG treatments (p < 0.0001). Likewise, SG 400 NV deviated from all other SG treatments (p = 0.0102, figure 2c).

This result suggests that [CO_2_] and vernalization act independently to influence expression of *FT. FT* displays independently regulated morning and evening expression peaks when exposed to warm-day, cool-night, and low red: far-red ratio as used here (Methods S1) (Song *et al*., 2015; Song *et al*., 2018). This could provide a mechanism for independent [CO_2_] and vernalization regulation of *FT*. We only harvested in the morning, and therefore subsequent experiments harvesting periodically across 24-h could clarify [CO_2_] and vernalization-specific *FT* regulation.

Like *FT, LFY* expression was uniform across genotypes and treatments with only two clear deviations. Importantly, one was elevated *LFY* in vernalized SG grown at 800 ppm [CO_2_] compared to other SG treatments (p < 0.0001; Fig. 2f). This is also a clear negative relationship between *FLC* and *LFY*, consistent with vernalization-induced early flowering at elevated [CO_2_]. We also observed elevated *LFY* in non-vernalized CG grown in 800 ppm [CO_2_] compared to other CG treatments (T_0_ to 3-wk *p* < 0.0001; Fig. 2e), consistent with our previous work (Springer *et al*., 2008), and associated with early senescence in this treatment. As *LFY* is associated with senescence (Ditt *et al*., 2011), this may also explain the high *LFY* expression at the last time point in non-vernalized SG in 800 ppm [CO_2_] (p < 0.0001; Fig. 2f). Additionally, although *FT* indirectly but positively regulates *LFY* (Blazquez & Weigel, 2000; Bouché *et al*., 2016), expression of *FT* and *LFY* were not positively correlated. Therefore, a mechanism other than *FT*, possibly *SOC1* (Blazquez & Weigel, 2000; Bouché *et al*., 2016), may induce *LFY* in these conditions. Like *LFY, SOC1* was elevated in vernalized SG grown at 800 ppm [CO_2_] compared to other SG treatments (p < 0.0001; Fig 2h). This result lends further support to this proposed mechanism of vernalization-induced early flowering at elevated [CO_2_] via changes in *SOC1* expression.

Our results demonstrate that repression of *FLC* via vernalization coupled with upregulation of *LFY* and SOC1 in vernalized SG grown in 800 ppm [CO_2_] restores early flowering as observed at current [CO_2_]. Our results also suggest that elevated [CO_2_] acts through other mechanisms to regulate floral integrator genes downstream of *FT*.

### Sequence variation at the *FLC* locus may underlie SG response

*FLC* expression correlates with delayed flowering in SG grown at elevated [CO_2_], and it is well known that *FLC cis* variation drives natural flowering time variation (Caicedo *et al*., 2004; Guo *et al*., 2015; Méndez-Vigo *et al*., 2016). Therefore, we compared the *FLC* sequences of CG and SG encompassing known *FLC* regulatory domains (Shindo *et al*., 2006; Liu *et al*., 2010; Marquardt *et al*., 2014). CG and SG varied relative to each other and to the *Arabidopsis* Columbia-0 (Col-0) reference (Fig. 3, Table S2), primarily within or near known sites of *FLC* regulation. Notably, variants that distinguished CG from SG occurred in the nucleation region, including the cold memory element (CME), and the vernalization response element (VRE), both within intron 1. Five single nucleotide polymorphisms (SNPs) occurred in the nucleation region, two unique to SG. Among these was an A to T substitution 4 bp upstream of the CME, which contains a 6-bp Sph/Ry motif necessary for *FLC* repression by vernalization (Yuan *et al*., 2016). Both CG and SG contain a substitution from A to C eight bp downstream from a second Sph/Ry motif which is 26 bp downstream of the CME and modulates *FLC* repression rate (Yuan *et al*., 2016). As the nucleation region is where H3K27me3 histone repressive marks begin to accumulate with vernalization (Pyo *et al*., 2014), these variants, alone or in combination, may alter the *FLC* repression rate.

**Figure 3.**
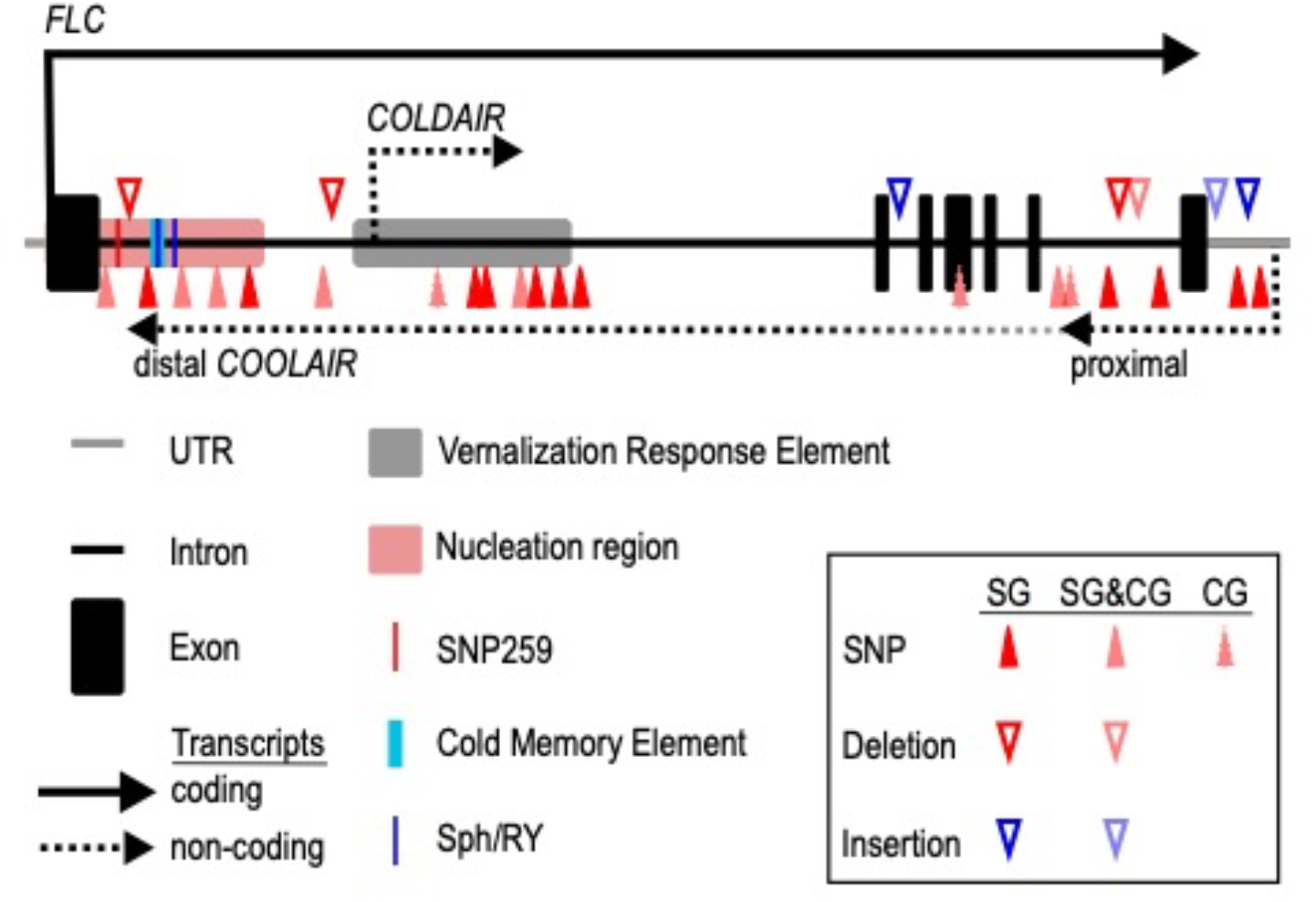
Selected Genotype (SG) displays sequence variation near known regulatory elements. relative to Columbia-0 (Col-0) and the Control Genotype (CG). *FLC* locus as determined from Col-0 accession (TAIR version 10) showing exons, introns, and known regulatory elements: SNP259 (Li et al. 2015 Genes & Dev.), Vernalization Response Element (Sung et al. 2006 Nat. Gen.), Nucleation Region (Angel et al. 2011 Nature), Cold Memory Element and the two Sph/Ry elements (Yuan et al. 2006 Nat. Gen.), and direction of *FLC* transcript and *COLDAIR* and *COOLAIR* non-coding RNAs. Colored arrow heads represent single nucleotide polymorphisms (SNP), insertions, or deletions in SG and CG relative to the Col-0 accession. Dark arrowheads occur in SG only, light arrowheads with solid boarder occur in SG and CG, light arrowheads with jagged boarder occur in CG only. All locations are approximate. See **Supplementary Table 2** for precise locations.

We identified multiple polymorphisms around the VRE, including a 6-bp deletion *c*. 38 bp upstream of the VRE, four SNPs within the VRE, and an additional SNP two bp downstream of the VRE, all unique to SG (Fig. 3, Table S2). The VRE modulates *FLC* repression through transcription of non-coding RNAs (ncRNA) *COLDAIR* and *COOLAIR*. These alter the rate at which Polycomb Repressive Complex 2 (PRC2) places histone repressive marks along *FLC* during vernalization (Marquardt *et al*., 2014; Pyo *et al*., 2014; Alamdar *et al*., 2019). Because transcription of *COLDAIR* begins within the VRE, these SNP variants may influence *COLDAIR* transcription to affect *FLC* repression contributing to CG and SG flowering differences.

Finally, SG contained a unique single-bp deletion 272 bp downstream of the *FLC* translational start site (Table S2). This deletion is near a known polymorphism called SNP259, containing a T to G substitution 259 bp downstream from the translational start site, occurring in accession Var2-6 from Sweden (Li *et al*., 2015). SNP259 alters the *FLC* vernalization repression rate by altering *COOLAIR* splicing (Li *et al*., 2015). Six additional polymorphisms distinguished SG from Col-0 and CG within the region spanning the truncated (proximal) *COOLAIR* variant (Fig 3., Table S2). Possibly, the deletion present in SG or the polymorphisms within the proximal region of *COOLAIR* alter *COOLAIR* splicing to influence *FLC* expression. More work is needed to determine whether these polymorphisms are responsible for the different genotype by [CO_2_] responses.

## Discussion

One possible effect of a future elevated [CO_2_] world is adaptation that occurs through altered flowering time. In Brassicaceae, this could occur through selection on standing variation of *FLC cis*-regulatory regions. While *FLC* is predominantly known for vernalization responsiveness (Searle *et al*., 2006), it also influences changes from perennial to annual life habits, drought response, and duration to seed germination (Chiang *et al*., 2009; Fletcher *et al*., 2016; Kiefer *et al*., 2017). As *FLC* function is similar across Brassicaceae species and genotypes (Kiefer *et al*., 2017), alteration of *FLC* expression may be a relatively rapid mechanism by which populations respond to rising [CO_2_].

Here we showed that altered *FLC* expression was causal in delaying flowering time at elevated [CO_2_] in an *Arabidopsis* genotype (SG) selected for high fitness under elevated [CO_2_], through direct manipulation in FLC expression via vernalization. Our study reveals a candidate mechanism, through *FLC*, that drives flowering time shifts in response to [CO_2_]. Further, we show that elevated [CO_2_] can induce vernalization sensitivity at 800 ppm that is not present at current CO_2_ (400 ppm) within a genotype, suggesting that large and unexpected responses in development, and flowering time more specifically, may occur in a high [CO_2_] world of the future.

We demonstrate that vernalization-induced *FLC* downregulation in SG grown at 800 ppm CO_2_ restores flowering time phenotypes that are characteristic of responses at 400 ppm CO_2_.

However, the specific mechanisms that caused altered expression of *FLC* at elevated CO_2_ are still unknown. One possibility is through [CO_2_]-induced changes in foliar sugars, as it is well known that elevated CO_2_ commonly increases plant carbohydrate status (Misra & Chen, 2015). For example, Col-0 produced more leaves and delayed flowering with increased growth media sucrose, similar to SG’s response to elevated [CO_2_] (Ohto *et al*., 2001). Several studies corroborate that sugar-related pathways alter *FLC* expression. For example, *sucrose non-fermenting-1 (Snf1)-related protein kinase 1* (*SnRK1*) subunit *KIN10* is involved in *FLC* repression (Jeong *et al*., 2015) and *Drosophila* polycomb-group proteins, analogous to *FLC* regulator PRC2, are modified post-translationally by sugar addition (Love & Mazer, 2021). In addition to overexpression of *FLC* and subsequent delayed flowering, this response was also coupled with a significant increase in sucrose in SG grown in elevated [CO_2_] (Springer *et*.*al*., 2008). Thus, foliar sugar modification due to [CO_2_] change is a potential mechanism of *FLC* epigenetic alteration.

Although *FLC* is a key determinant of flowering time, the similar alteration of *LFY and SOC1* here and in previous work (Springer *et al*., 2008) suggests that elevated [CO_2_] acts broadly to influence flowering gene regulation. Possibly due to the timing of tissue harvest, we did not observe predicted increases in *FT* expression in response to elevated [CO_2_]. However, we know from previous work that *SOC1* is positively regulated by *FT* within the photoperiod and sugar sensing pathways. In addition to being downstream of *FLC, LFY* functions within the senescence and sugar sensing pathways, both of which can explain *LFY’s* unexpected upregulation in non-vernalized elevated [CO_2_] treatments. Like SOC1, *LFY* expression is regulated by the key indicator of carbohydrate levels, *T6P* (trehalose-6-phospate) (Dijken *et al*., 2004; Wahl *et al*., 2013) and by the carbohydrate-sensitive transcription factor *SQUAMOSA BINDING PROTEIN LIKE 3* (*SPL3*) (Golembeski *et al*., 2014; Bouché *et al*., 2016). It is possible that *LFY’s* response to elevated [CO_2_], perhaps even independently of SOC1, and, therefore, to elevated carbohydrate status, may be promoting senescence in the already very early flowering CG genotype consistent with previous studies (Ditt *et al*., 2011). *LFY* may, along with *FLC*, be contributing to both early flowering and senescence in vernalized SG grown at elevated [CO_2_].

Our work highlights a mechanism through which rising [CO_2_] may influence flowering time, specifically through sustained high levels of *FLC*, potentially modulated by vernalization. However, extreme weather events and heat-waves (Allen *et al*., 2018), may cause devernalization, leading to *FLC* upregulation even after stable repression (Bouché *et al*., 2015). We also show that [CO_2_] may act through *FT* and *LFY* as well as *FLC*. Therefore, interplay of [CO_2_] and other climate change factors, on flowering is likely highly complex, warranting further exploration.

## Supporting information

Supplemental Figure 1, Supplemental Table 1, Supplemental Table 2, Methods, and will be used for the link to the file on the preprint site.

## Acknowledgements

The authors thank Drs. Carrie Wessinger, Clint Springer, John Kelly, Ms. Rebecca Orozco, and the University of Kansas Genome Core Facility as this work would not have been possible without their initial discussions, findings, plant materials, and technical support. We thank Upendra Kumar Devisetty at Cyverse for their assistance with creation of blastn_custome_outfmt6_subject_sequence-2.6.0 application. CyVerse is supported by the NSF under Award Numbers DBI-0735191, DBI-1265383, and DBI-1743442. This work was supported by NSF (IOS-1457236) to JKW and LCH, the NIH Research and Academic Career Development Award (IRADCA) fellowship K12GM063651 to HK-S, and the University of Kansas Department of Ecology and Evolutionary Biology and Botany Endowment Fund for support of ALH-C. This paper is dedicated to University of Kansas alumni of the Ward Laboratory.

## Author contributions

ALH-C, HK-S, LH, and JKW designed the experiments. ALH-C performed the experiments. ALH-C and HK-S analyzed the data. ALH-C, HK-S, LH and JKW wrote the manuscript. LH and JKW edited the manuscript. All authors approved of the manuscript.

## Data availability

The data that supports the findings of this study are available within the paper and its Supporting Information.

