## Supplemental Figure 1, Supplemental Table 1, Supplemental Table 2, Methods, and will be used for the link to the file on the preprint site. for "*FLOWERING LOCUS C* drives delayed flowering in *Arabidopsis* grown and selected at elevated CO_2_"

**Annals of Botany Supporting Information**

The following Supporting Information is available for this article:

**Figure S1:** Biomass at bolt

**Table S1:** Primer sequences

**Table S2:** Columbia-0, CG, SG *FLC* polymorphism locations

**Methods S1:** Materials and methods details

**Fig. S1** Biomass at bolt

**Fig. 1 Biomass (mg) at bolt of *Arabidopsis thaliana* Control Genotype (CG) and Selected Genotype (SG)** in non-vernalized (NV) and vernalized (V) conditions under elevated [$CO_2$] (800ppm) and 'current' [$CO_2$] (400ppm). Boxes indicated 25% and 75% quartiles and the heavy black line represents the median. Whiskers extend to highest value with 1.5 x the interquartile range. Lower-case letters indicate differences in significance. Each point represents a single individual.

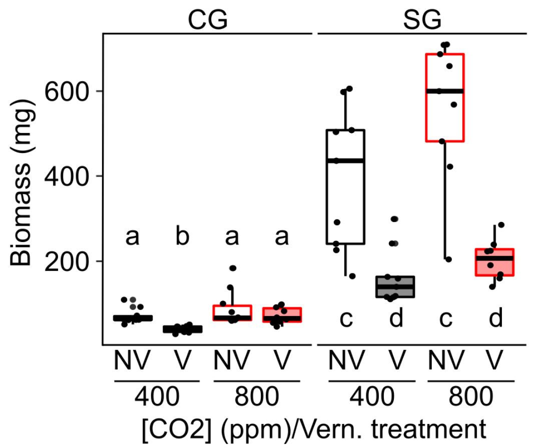

**Table S1** Primer sequences

| **Primer name** | **Sequence** | **Source** |
|---|---|---|
| Q_FLC_Splc_F | GCCAAGAAGACCGAACTCATGTTGA | HK-S, primer designing tool – NCBI, (Piriyapongsa *et al.*, 2009) |
| Q_FLC_Splc-R | CAACCGCCGATTTAAGGTGGCTA | HK-S, primer designing tool – NCBI, (Piriyapongsa *et al.*, 2009) |
| QC_AT5G08290_F | GGATGAGACCTGTATGCAGATGGATGAG | (Hong *et al.*, 2010) |
| QC_AT5G08290_R | GAGAGCCCAGTTGATCTTGTTGTTGTTAC | (Hong *et al.*, 2010) |
| QC_AT4G27960_F_UBC9 | GCTCTCACAATTTCCAAGGTGCTGC | (Hong *et al.*, 2010) |
| QC_AT4G27960_R_UBC9 | AGGGCTCTTCCTTAAGGACAGTATTTGTG | (Hong *et al.*, 2010) |
| Q_LEAFY_F | CGGCGAAGATAGCGGAGTTA | ALH-C, primer designing tool-NCBI, (Piriyapongsa *et al.*, 2009) |
| Q_LEAFY_R | AGAGCGTGATGAGTACCGGA | ALH-C, primer designing tool-NCBI, (Piriyapongsa *et al.*, 2009) |
| Q_FT_F | AGCATAGCTCAAACATGTTGCT | ALH-C, primer designing tool-NCBI, (Piriyapongsa *et al.*, 2009) |
| Q_FT_R | TAGCTTGGGTGTGGGCTTTT | ALH-C, primer designing tool-NCBI, (Piriyapongsa *et al.*, 2009) |
| Q_SOC1_F | TCAAGTCGAGAGACTAGAGGTC | ALH-C, primer designing tool-NCBI, (Piriyapongsa *et al.*, 2009) |
| Q_SOC1_R | TGTAGAGTTGCCACCGTTGT | ALH-C, primer designing tool-NCBI, (Piriyapongsa *et al.*, 2009) |

**Table S2** Columbia-0, CG, SG *FLC* polymorphism locations

| Position from ATG | Col | CG | SG | Location type |
|---|---|---|---|---|
| -1813 | g | t | g | IG |
| -1783 | a | a | g | IG |
| -1767 | c | c | t | IG |
| -1671 | t | t | g | IG |
| -1668 | t | t | g | IG |
| -1383 | a | a | - | IG |
| -1382 | t | a | - | IG |
| -1373 | t | t | c | IG |
| -1168 | g | g | t | IG |
| -1046 | t | c | t | IG |
| -1002 | g | g | a | IG |
| -736 | a | - | - | IG |
| -735 | a | c | a | IG |
| -309 | c | a | a | IG |
| -292 | g | g | a | IG |
| -230 | a | g | g | IG |
| -177 | a | - | - | IG |
| -176 | a | a | c | IG |
| 0 | a | a | a | ATG |
| 1 | t | t | t | ATG |
| 2 | g | g | g | ATG |
| 216 | g | a | a | I1,NR |
| 259 | g | g | g | I1,NR,SNP259 |
| 272 | a | a | - | I1,NR |
| 466 | a | a | t | I1,NR |
| 470 | t | t | t | I1,NR,CME start |
| 473 | t | t | t | I1,NR,CME,Sph/RY-1 |
| 506 | t | t | t | I1,NR,Sph/RY-2 |
| 514 | a | c | c | I1,NR |
| 719 | a | g | g | I1,NR |
| 884 | t | t | g | I1,NR |
| 1280 | a | g | g | I1 |
| 1290 | c | c | - | I1 |
| 1291 | a | a | - | I1 |
| 1292 | a | a | - | I1 |
| 1293 | a | a | - | I1 |
| 1294 | g | g | - | I1 |
| 1295 | g | g | - | I1 |
| 2228 | a | t | a | I1,VRE,*COLDAIR* |
| 2447 | a | a | g | I1,VRE,*COLDAIR* |
| 2450 | c | c | t | I1,VRE,*COLDAIR* |
| 2765 | g | t | t | I1,VRE |

| Location Key | |
|---|---|
| SNP259 | G to T substitution alters response |
| VRE | Vernalization Response Element |
| NR | Nucleation Region |
| CME | Cold Memory Element |
| *COOLAIR* | *COOLAIR* |
| *COLDAIR* | *COLDAIR* |
| E# | E1 = Exon 1 |
| I# | I1 = Intron 1 |
| Sph/RY-# | Motif-# position from gene start |

| Sources used to determine locations | |
|---|---|
| *FLC* | TAIR AT5G10150 (from 2017)<br>ATG at TAIR position 3179360<br>(Huala *et al.*, 2001) |
| SNP259 | (Li *et al.*, 2015) |
| VRE | (Sung *et al.*, 2006) |
| NR | (Angel *et al.*, 2011) |
| CME, Sph/RY | (Yuan *et al.*, 2016) |
| *COOLAIR* | (Marquardt *et al.*, 2014)<br>GenBank: HG975389.1 |
| *COLDAIR* | (Heo & Sung, 2011)<br>GenBank: HG975388.1 |

| | | | | |
|---|---|---|---|---|
| 2790 | a | a | c | I1,VRE |
| 2869 | g | g | a | I1,VRE |
| 2946 | g | g | a | I1 |
| 3899 | c | c | c | I2 |
| | - | - | t | I2 |
| | - | - | t | I2 |
| 3900 | c | c | c | I2 |
| 4129 | c | t | c | E4 |
| 4801 | t | a | a | I6 |
| 4864 | a | g | a | I6 |
| 5127 | a | a | g | I6 |
| 5135 | a | a | - | I6 |
| 5212 | g | - | - | I6 |
| 5396 | c | c | a | I6 |
| 5669 | a | a | a | I6 |
| | - | a | a | 3'UTR,*COOLAIR* Prox.* |
| | - | g | g | 3'UTR,*COOLAIR* Prox.* |
| 5670 | a | a | a | 3'UTR,*COOLAIR* Prox.* |
| 5791 | c | c | t | 3'UTR,*COOLAIR* Prox.* |
| 5806 | c | c | c | 3'UTR,*COOLAIR* Prox.* |
| | - | - | t | 3'UTR,*COOLAIR* Prox.* |
| 5807 | a | a | a | 3'UTR,*COOLAIR* Prox.* |
| 5843 | a | a | t | *COOLAIR* Prox.* |

Positions based on Col-0 sequence (TAIR). **COOLAIR* has several splice variants (Li *et al.*, 2015). Represented is the approximate location of the *COOLAIR* Proximal (Prox.), or truncated, variants. Translational start site blocked in grey for reference. No position number indicates an insert in either SG or CG.

**Methods S1** Materials and Methods

**Plant materials and growth conditions**

*Arabidopsis thaliana* (L.) Heynh SG and CG genotypes (Ward *et al.*, 2000; Springer *et al.*, 2008) were grown from seed in 500 ml pots filled with Berger BM6 Blend soil. Seeds were cold stratified at 4°C for five days to promote uniform germination. Plants were grown in growth chambers (Conviron BDR16, Winnipeg, Canada), maintained at 400 ppm ('current') and 800 ppm ('elevated') [$CO_2$], hereafter e[$CO_2$], 22:18°C (day:night), and 60:80% relative humidity. Plants were grown in long day conditions, 14:10 h photoperiod, *c.* 650 µmol $m^{-2}$ $s^{-1}$ light intensity, using Phillips Metal Halide (Phillips Lighting Company, China) and Phillips EcoAdvantage (Hg) incandescent bulbs (Phillips Lighting, Mexico). Incandescent bulbs lowered the far-red light ratio, making the spectrum quality more consistent with natural light (Darko *et al.*, 2014).

Plants were watered twice daily and given half-strength Hoagland's solution once daily. When plants developed four to five true leaves, a subset (*n*=20) of SG and CG were moved into separate chambers programmed for vernalization (5°C) at both current and e[$CO_2$]. To maintain growth-chamber function in the vernalization plus high light treatment, in consultation with the manufacturer, over-head lights were shut off and the temperature increased from 5°C to 10°C for three ten-minute intervals during the light period. SG and CG plants were vernalized for four consecutive weeks, then returned to current or e[$CO_2$] to record flowering. Rosette leaf number and shoot weight were recorded when the meristem reached 1 cm.

**Harvesting samples**

Whole rosettes were harvested from vernalized and non-vernalized SG and CG plants for gene expression. Whole rosettes from 3 to 4 individual plants were harvested between dawn and noon at five time points: vernalization start (time 0), then two, three, and four weeks after the start of vernalization, and five days after vernalization ended. Tissue was stored at -80°C until processed, cooled in Liquid N, and ground using a Bead Bug 6 Microtube Homogenizer (Benchmark, China). Powdered tissue samples were stored at -80°C in 0.5 ml SureSeal S Microtubes (Midwest Scientific, USA) for RNA isolation.

**RNA isolation and quantitative RT-PCR**

RNA was isolated using the Illustra RNAspin Mini kit with on-column DNAse treatment (GE Healthcare, Fisher Scientific, Germany), concentrations were determined using a NanoDrop™ 1000 ND Spectrophotometer (Thermo Fisher Scientific, USA), two micrograms of RNA was used for cDNA synthesis using the iScript™ cDNA Synthesis Kit (Bio-Rad, Hercules, CA, USA). Amplification of two or three technical replicates of each biological replicate samples (*n* = 3) was monitored in real time using SYBR Select Master Mix (Applied Biosystems, Fisher Scientific, USA) on a QuantStudio™ Real-Time PCR system (Thermo Fisher, USA). Expression of *FLOWERING LOCUS C* (*FLC*), *FLOWERING LOCUS T* (*FT*), and *LEAFY* (*LFY*) transcript levels were assessed and normalized relative to reference genes UBC9 (AT4G27960) and YLS8 (AT5G08290) (Czechowski *et al.*, 2005; Hong *et al.*, 2010) using the DCT method. Target gene expression levels were calculated by using the following excel formula: = 2^(average(GOR ) – (average

(GOI)) as described (Ito *et al.*, 2012; Kinmonth-Schultz *et al.*, 2021). Primer sequences are in **Table S1**.

**Sequencing *FLC***

The *FLC* sequence was pulled from assembled genomes for SG and CG and sequenced through the University of Kansas Genomics CORE facility. Specifically, gDNA was isolated using the DNeasy Plant kit (Qiagen, Denmark) from two pooled, fully inbred plants from both the SG and CG lines, the libraries were prepared using the TruSeq DNA PCR-Free kit and sequenced on the HiSeq RR-PE100 system (Illumina, USA). This resulted in approximately 188 million reads in total or about 94 million reads per pooled sample, and about 200x coverage per genotype. These 100-bp reads were assembled using the CLC Genomics Workbench (CLC Genomics Workbench (RRID:SCR_011853) against the default *Arabidopsis thaliana* reference genome (Col-0) downloaded from The Arabidopsis Information Resource (TAIR version 10) (Rhee *et al.*, 2003). The *FLC* sequences were pulled from the SG and CG genome sequences using the Discovery Environment in Cyverse (www.cyverse.org) (Polański *et al.*, 2018) using the following applications: Create_BLAST_database-2.6.0+ to create a database from each genome sequence and blastn_custom_outfmt6_subject_sequence-2.6.0+ to extract the *FLC* sequence from each genome database using Col-0 *FLC* sequence (TAIR) (Rhee *et al.*, 2003) as the query. The sequences were then aligned using Multiple Sequence Alignment (MUSCLE) (Edgar, 2004) and visually compared for differences between SG, CG, and the Columbia-0 reference sequence in a Plasmid Editor (aPE)(Heathward *et al.*, 2016).

**Statistical analyses**

Leaf number and gene expression were analyzed across genotypes, [$CO_2$], and vernalization treatments using ANOVA and Tukey post-hoc test (*aov*, *TUKEYHSD* R version 4.11). To compare gene expression of *FLC*, *FT*, and *LFY* over time, we grouped the plots by [$CO_2$] and vernalization treatment and tested each genotype separately.
